# Uncovering the geometry of color space with magnetoencephalography (MEG)

**DOI:** 10.1101/2020.08.10.245324

**Authors:** Isabelle Rosenthal, Shridhar Singh, Katherine Hermann, Dimitrios Pantazis, Bevil R. Conway

## Abstract

The geometry that describes the relationship among colors is unsettled despite centuries of study. Here we present a new approach, using multivariate analyses of direct measurements of brain activity obtained with magnetoencephalography to reverse-engineer the geometry of the neural representation of color space. The analyses depend upon determining similarity relationships among the neural responses to different colors and assessing how these relationships change in time. To evaluate the approach, we relate patterns of neural activity to universal patterns in color naming. Control experiments showed that responses to color words could not decode activity elicited by color stimuli. The results suggest that three patterns of color naming can be accounted for by decoding the similarity relationships in the neural representation of color: the association of warm colors such as reds and oranges with “light” and cool colors such as blues and greens with “dark”; the greater precision among all languages in naming warm colors compared to cool colors; and the preeminence of red.

---

It is popular to depict color in three-dimensional spaces in which distances between colors correspond to perceptual similarity [1]. But these geometric representations are cartoons that do not accurately capture a uniform representation of color [2, 3]. Many attempts have been made to develop a truly perceptually uniform color space, including Munsell [4], CIELUV/CIELAB/CIECAM02 [5], the Natural Color System [6], and the Optical Society of America Color System [7]. Each has a unique 3-dimensional geometry obtained from fitting models to perceptual judgements. But none of these, or any other color space developed to date, is accurate or formally defined [2, 8]. Moreover, interpretation of the perceptual judgments, and by extension the relationship among colors that those judgements reveal, may be complicated by confounds introduced by task demands [9].

The color percept associated with a stimulus can be decoded from functional magnetic resonance imaging activity [10], MEG and EEG in humans and monkeys [11-14], or responses of single neurons in monkeys [15, 16]. Moreover, as many of these studies show, the extent to which activity patterns reflect the sequence of hues in the color wheel can be used to identify neurons and brain areas that are likely involved in encoding color. In such experiments, the sequence of hues reflects the crude common property of all color spaces but does not capture the geometry of color space. In the experiments presented here the logic is flipped: we take as the central problem gaps in knowledge about the geometry of the neural representation of color, and aim to reverse-engineer this geometry from direct measurements of brain responses obtained with magnetoencephalography (MEG). We take up this aim using a decoding approach [17, 18]. Our goal is to determine the neural geometry of color by assessing the similarity relationships among the patterns of MEG activity elicited by a set of colors: colors with more similar patterns of MEG activity are considered more similar. This approach is analogous to the way perceptual similarities among colors are established using asymmetric color matches [8], where participants might be asked to pick a color that is most similar, but not identical, to the target. We use stimuli with equal cone contrast, defined by the cone-opponent axes that the retina uses to encode color (**Figure 1a**) [19-22]. Because MEG is thought to predominantly reflect cortical responses, the experiments reveal the transformation of retinal signals by the cortex.

**Figure 1.**
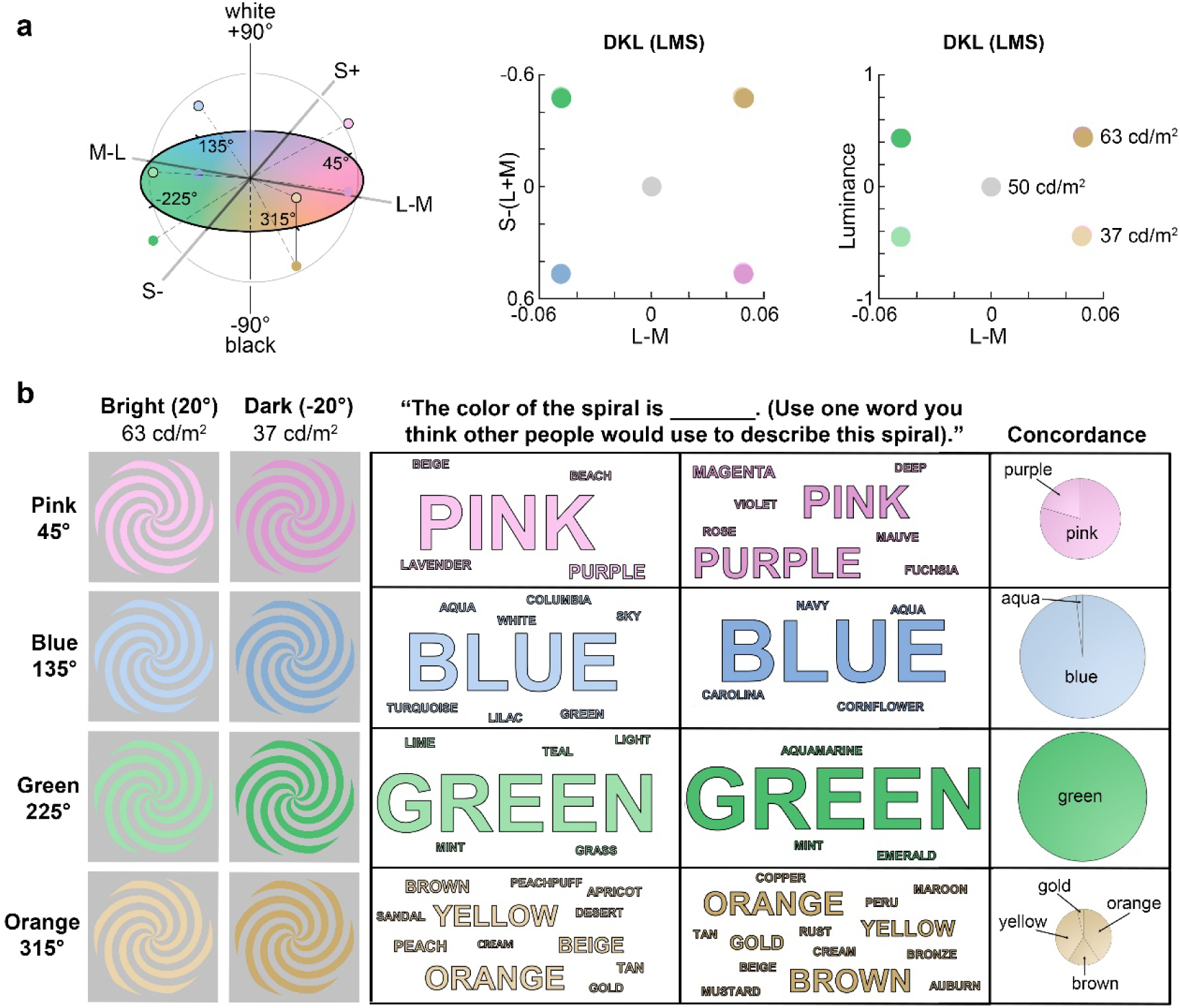
Specification of the stimulus colors, and how they are named. **a**, Stimuli were colored spirals surrounded by neutral adapting gray; the colors defined within a color space defined by the cardinal cone-opponent mechanisms, indicated by the right panel [19, 20]. The left panels show the cone and luminance contrast of the stimuli, relative to the gray adapting background. **b, Left:** Configuration of the stimuli; spirals subtended 10° of visual angle, and varied in hue (rows) and luminance (columns), as defined in DKL color space. **Center:** Word clouds, reflecting the distribution of color names assigned to the stimuli. Font size reflects how many subjects assigned that word to the stimulus (N=70). The smallest words (ex: ‘lavender’) represent just one response; the largest words (‘green’ and ‘blue’ in dark hues column) represent 66 responses. Subjects were mTurk volunteers (N=52) and the participants in the main MEG experiment (N=18). **Right:** Concordance, measuring how similar responses were within hues and across luminance sets (N=70). The size of the pie chart reflects the proportion of participants who used the same word across luminance sets, and the content of chart indicates which words were used for both luminance sets. Almost all people used “green” for both high and low luminance contrast versions of that hue (large pie for hue 225°), whereas relatively few people used the same term for both high and low luminance contrast versions of the yellow/brown/ochre stimulus (small pie for color 315°), and among these people, several terms were used (many pie slices for 315°). The same term was used for both versions of the given hue (“concordance” across luminance sets) for: pink= 55.6% of participants; Blue=85.7%; Green=88.6%; Orange=38.6%. The most commonly used term for hue 135° (blue) at the light|dark luminance level was used by 87%|94% of participants; the most commonly used term for hue 225° (green) was used by 90%|94% of participants. For hue 45° (pink), there was lower consensus (81%|50%) and lower concordance (44% used pink for light and dark versions; 11% used purple). For hue 315° (orange), there was no clear consensus: the most commonly used term, orange, was only used by 30% of participants (at both luminance levels). There was no difference in the level of concordance between blue and green (chi-square test of proportions, p=0.61; Bonferroni alpha = 0.0083); similarly, there was no difference in concordance between pink and orange (p=0.04; Bonferroni alpha = 0.0083).

To evaluate the approach, we relate the patterns of neural activity elicited by different colors to universal patterns in color naming, guided by the idea that the cross-linguistic patterns in color naming derive from an underlying geometry of the neural representation. We consider three universal aspects of color naming: First, warm colors such as reds and oranges are associated with “light” and cool colors such as blues and greens are associated with “dark” [23]. Second, among all languages there is a greater precision in naming warm colors compared to cool colors [24, 25]. And third, after black and white, red is the most important basic color term [26].

## Results

We measured MEG responses to 8 colored spirals; the stimuli comprised four hues at two luminance-contrast levels; the cone contrast was equated for all the stimuli, and all stimuli modulated activity in all three cone types (**Figure 1a**). We also measured responses to achromatic text of the words “green” and “blue”. The experimental paradigm represents a compromise between having a large enough representation of color space, a set of colors with well-defined and balanced cone activation, and a small enough set of stimuli to enable each stimulus to be presented a large number of times to ensure sufficient power for decoding analysis. Except for the words, the stimuli had the same shape and texture, ensuring the same pattern of retinotopic activation; thus the extent to which patterns of activation elicited by one spiral versus another could be decoded can only be attributed to differences in color. The stimuli consisted of two warm colors (colors elicited by L>M cone activity; pink, orange) and two cool colors (blue, green). Warm colors are associated with “light”, and cool colors (blues, greens) are associated with “dark” among all languages, even those with the most limited numbers of consensus color terms [27]. Among the 11 basic color categories (red, pink, orange, yellow, brown, green, blue, purple), the cool colors (green, blue) are not distinguished from each other by differences in lightness, whereas the warm colors are [23-25, 28]. Consistent with this universal pattern, participants often used different terms for the light and dark versions of warm colors and the same term for light and dark versions of the cool colors (**Figure 1b**).

### Classifying colors using MEG responses

MEG activity elicited by the colored spirals was measured in 18 people (all of whom also participated in the color-naming experiment). Each spiral was presented 500 times—a large number of trials, intended to ensure sufficient power for the decoding analyses. For each of the 18 participants, a classifier was trained to decode stimulus color from the pattern of MEG results. The classifiers accurately identified the colors of the spirals based on MEG data, using separate data to train and test the classifiers (**Figure 2a**). The classifier trained to decode which among 8 colors gave rise to the pattern of MEG activity reached significance at 55 ms (95% CI: [50,55]), with a peak decoding time at 115 ms (95% CI: [105,125]). Across subjects, the time to peak decoding ranged between 95-260 ms. The performance of these classifiers cannot be attributed to confounds introduced by eye position or pupil diameter (data not shown). Pink was decodable with the shortest latency (for light stimuli, pink 70 ms [65, 70, 95 % C.I.]; blue 80 ms [80 80], green 80 ms [80 80], orange 80 ms [80 80]; for dark stimuli, pink 55 ms [55, 60]; blue 75 ms [75 90], green 75 ms [75 75], orange 75 ms [75 75]. Moreover, peak decoding accuracy differed for each of the eight colors (Friedman test, p=0.009, **Figure 2b**). Among all the colors, pink (regardless of luminance contrast) showed the highest decoding accuracy (Wilcoxon signed rank test, light stimuli, pink>blue, p=0.004; pink>green, p=0.001; pink>orange, p=0.02; dark stimuli, pink > blue, p=0.16; pink>green, p=0.22; pink > orange, p=0.013).

**Figure 2.**
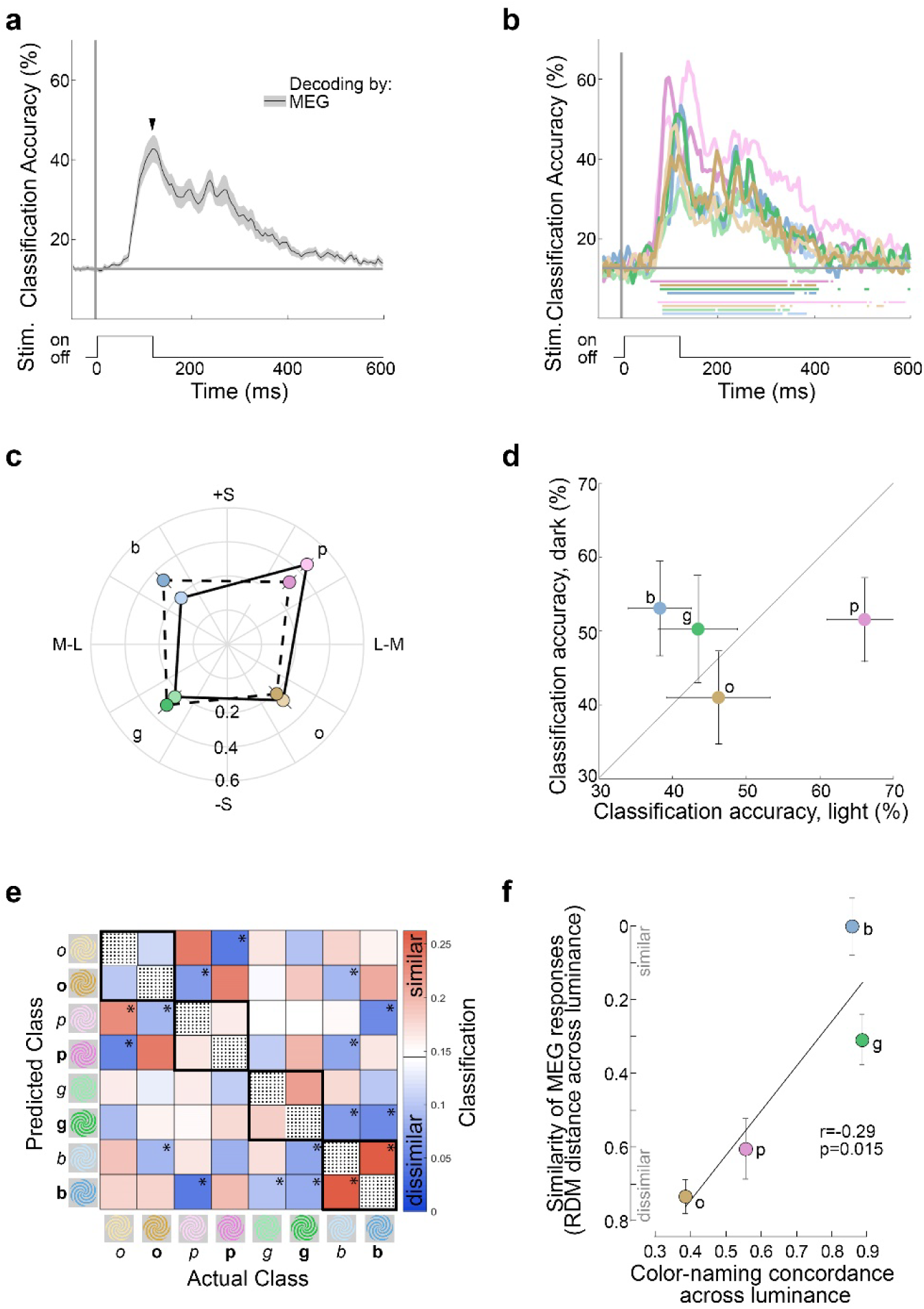
Decoding stimulus color from MEG data. **a**, Average classification accuracies across time for decoding the color of the 8 spirals, averaged over all 8 classification problems (N=18 participants; chance = 1/8; shading shows 95% C.I.). Step at the bottom shows the stimulus duration. The classifier was trained and tested for each subject individually using the 25 MEG sensors most selective for stimulus class on ANOVA. 0ms on the x-axis represents stimulus onset; the classifier was trained and tested in bins of 5ms. The horizontal line shows the times at which the classifier guessed the stimulus color above chance (bootstrapped 1000 times and FDR-corrected, p<0.05). **b**, Average classification accuracy for decoding each of the 8 colors (line color corresponds to stimulus color); other conventions as for panel (a). **c**, The peak decoding accuracy of each color, plotted in DKL hue angle space (b, blue; p, pink; o, orange; g, green). Color accuracies were sampled from each subject at their overall individual peak decoding time, and then averaged across subjects. Radius = averaged accuracy across individual classifiers. Error bars = SEM. **d**, Classification accuracy for light colors versus dark colors. Values and error bars as in (c); dark versions of cool colors showed higher classification accuracy compared to light versions, and light versions of warm colors showed higher classification accuracy compared to dark versions (repeated measures two-way ANOVA, interaction between wamrth and luminance, p=0.0116). **e**, Representational dissimilarity matrix (RDM) showing the classification forced error, at each subject’s individual peak decoding time, averaged across subjects. The heatmap shows the probability, normalized by column to sum to 1, that the classifier would assign a given spiral identity (vertical axis) when tested with data obtained by presenting a given spiral (horizontal axis). The classifier was trained in the same manner as in (a), but forced to guess the most likely spiral besides the correct option. White on the color bar = chance level. Asterisks indicate that the value in a box is either significantly above chance (red boxes) or below chance (blue boxes) (bootstrapped 95% confidence intervals; n=1000). **f**, RDM distances between warm colors and cool colors within each luminance level (p = 0.02).

At the time point of peak classification accuracy (determined for each participant, across all eight colors), classification accuracy varied by hue (repeated measures two-way ANOVA on rank transformed data, p=0.0387) but not by luminance level (p=0.8929)(**Figure 2c**). The variance in classification accuracy by hue was not predicted by a simple asymmetry in classifying colors along the cardinal cone-opponent (L-vs-M or S axes, **Figure 2c**). Classification was comparable for colors that vary along the L-M axis: warm colors (defined as L-M; pink, orange) compared to cool colors (M-L; green, blue) (Wilcoxon signed-rank test, N=18, p= 0.2485); and classification was comparable colors that vary along the S axis: S-increment colors (pink, blue) compared to S-decrement colors (orange, green) (p=0.0741). Instead the variance in classification accuracy by hue was accounted for by two asymmetries. First, classification was slightly worse for colors corresponding to the daylight locus (orange-blue) compared to the anti-daylight locus (pink-green) (Wilcoxon signed-rank test, p=0.0108). Second, classification accuracy co-varied with warm-cool and luminance contrast, such that for warm colors, the light versions were more accurately classified compared to the dark versions, while for cool colors, the dark versions were more accurately classified compared to the light versions (**Figure 2d**, repeated measures two-way ANOVA, no main effects, significant interaction between warm-cool and luminance contrast, p = 0.0116). This result provides a neural correlate of the association of warm colors with higher lightness, and cool colors with lower lightness which is evident in the universal pattern in color naming across languages [23].

### Representational similarity

We sought to analyze the MEG results in a way that was analogous to the methods used to infer the geometry of perceptual color space. Data from psychophysics experiments that are the basis for knowledge of the geometry of color space depend on an evaluation of the similarity relationships among colors. In practical terms, these relationships can be measured through an asymmetric matching paradigm, in which participants are asked to identify among a set of colors the color that is most similar, but not identical, to a given sample. We sought to apply this logic to analyze the MEG data to determine the geometry of the neural representation of color by asking: what color does the classifier pick when forced to pick any color except the correct color? In other words, what color besides the test color elicits a neural representation that is most similar to the neural representation of the test color? The pattern of errors (**Figure 2e**) provides an estimate of the similarity in the neural representation among colors that is analogous to the similarity among colors recovered in psychophysical color-ordering tasks. Colors that show more similar neural response patterns correspond to a closer relationship in the geometry of color space. (We also performed an analysis on the errors of the classifier performance, but the number of errors is relatively low compared to the number of trials; the forced-error analysis has more power and is more directly analogous to asymmetric color matching paradigms used in psychophysics to evaluate similarity relationships among colors.)

The heatmap shown in **Figure 2e** is largely symmetric about the identity diagonal, confirming that the pattern of results is meaningful (Pearson correlation coefficient comparing the two halves of the matrix, r=0.9639, p=2E-16). For example, the confusion between colors of the same hue but different luminance were almost identical: Dark orange was predicted when light orange was the actual color with almost the same likelihood that light orange was predicted when dark orange was the actual color, and so on for the four hues (see each of the four black-outlined squares along the inverse diagonal). But the confusion between stimuli of the same hue but different luminance varied among the four hues. For example, the pattern of MEG activity for blue was similar for light and dark versions of blue, while the pattern of MEG activity for orange was dissimilar for light and dark versions of orange. Rank ordered by hue, the similarity relationships corresponded to the concordance rates for color naming (**Figure 2f;** correlation coefficient = −0.2848, p=0.0153), and the distance between hues across luminance contrast was greater for warm colors (Kruskal-Wallis test, p=0.0034). The results show that the neural representation of warm colors is more impacted by luminance contrast than the neural representation of cool colors, which is consistent with the color-naming patterns shown in **Figure 1b**, in which blue and green at different luminance levels were typically assigned the same labels while pink and orange at different luminance levels were often given different labels.

**Figure 3** quantifies the similarity relationships among the colors. **Figure 3a** shows multidimensional scaling plots constructed from confusion matrices to generate 2-dimensional pictures of the relationship among the colors—the MDS plots are not simply a replotting of the confusion matrices, but instead show the best spatial representation of the RDMs as constrained by two orthogonal dimensions (constraining to two dimensions is warranted given the stimuli were varied along two dimensions, hue and luminance contrast). The MDS plots were produced using the stress metric. In the MDS plots, data points that are closer together correspond to neural representations that are more similar, i.e. more likely to be mistaken if the classifier fails. Each MDS plot shows the similarity relationships among colors for a single snapshot in time. The RDM data used to produce the MDS plots were sampled relative to the time to peak decoding for each subject, with negative values corresponding to times before peak decoding was achieved. **Figure 3a** shows the MDS plots at six time points, three spanning times around the peak decoding of stimulus onset (−5 ms, 0 ms, 5 ms) and three corresponding to times around the peak decoding of stimulus cessation (105 ms, 110 ms, 115 ms). The goodness of fit ranges of the six MDS plots to the measured differences: R=0.835307 – 0.951547 and p=7.84e-15 – 3.21e-8 (see Figure 3 legend). The MDS plots show qualitatively two striking patterns: first, each MDS plot can be oriented such that the light and dark hues are displaced vertically, and in this configuration the hues at both luminance levels for all six MDS plots follow a similar order: cool color, warm color, warm color, cool color. Second, the MDS plots corresponding to peak decoding of stimulus cessation are distinguished by a dramatic separation of the hues by luminance contrast.

**Figure 3.**
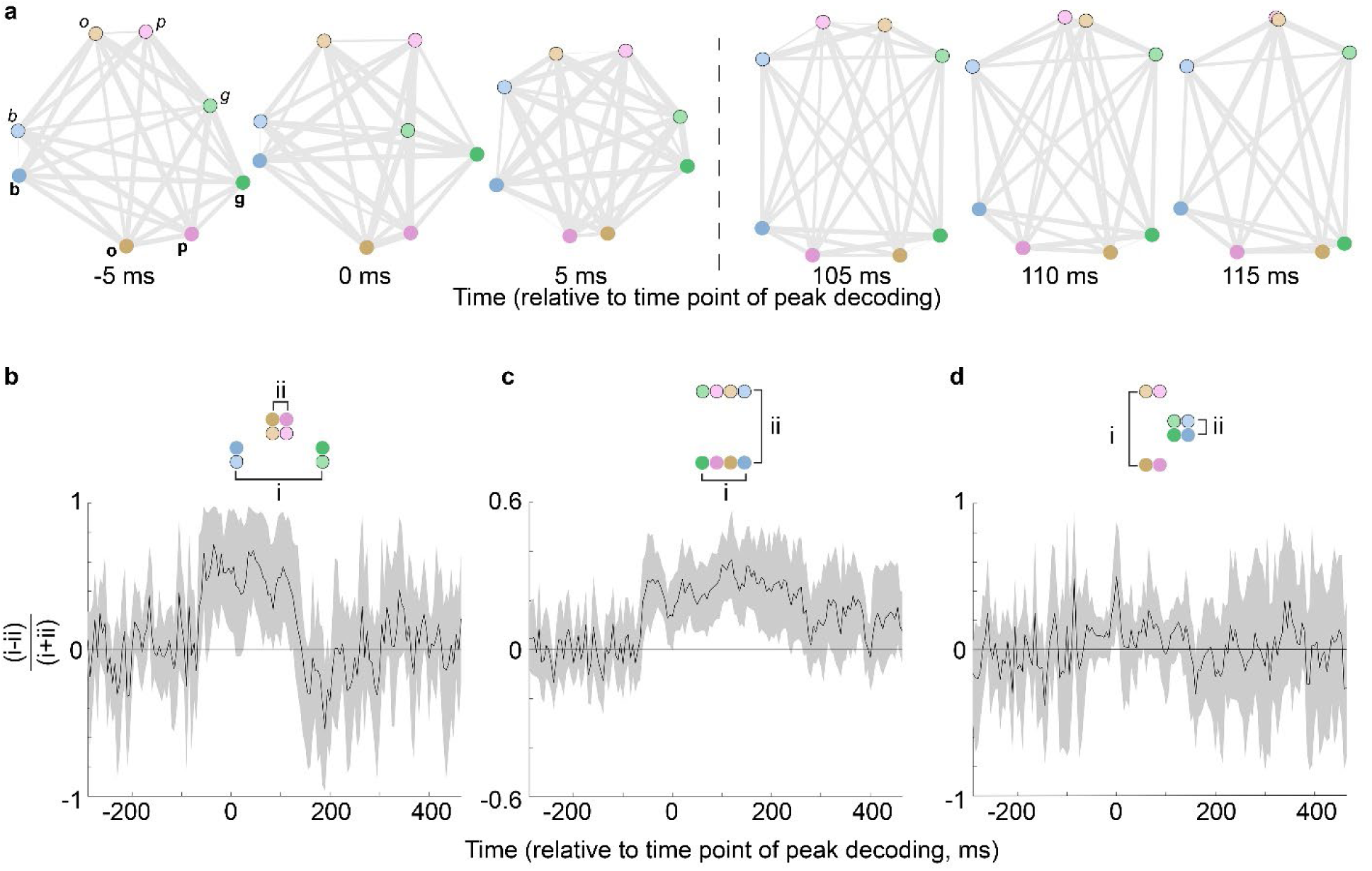
Multidimensional scaling (MDS) representation of the similarity relationships among colors determined by MEG. **a**, Six MDS plots sampled from different time points relative to the time point of peak decoding accuracy 0 ms corresponds to peak decoding of stimulus onset; 110 ms corresponds to peak decoding of stimulus cessation; the vertical dashed line separates MDS plots corresponding to onset from those corresponding to stimulus cessation. Symbol color corresponds to the color of the stimulus, and the light versions are outlined in black to further distinguish the data points. Shorter distances correspond to higher levels of classification confusion (higher similarity). The area of each line is directly proportional to the dissimilarity in the representation of the colors joined by the line. Hues are clearly separated by luminance contrast (light colors are at the top in each panel); and arranged by hue in a sequence, from left to right, of blue, orange (or pink), pink (or orange), green. Fit of MDS for each panel, left to right: Pearson’s; R=0.84, p=2×10^−8^; R=0.84, p=3×10^−8^; R=0.85, p=2×10^−8^; R=0.91, p=1×10^−11^; R=0.94, p=9×10^−14^; R=0.95, p=7×10^−15^. **b**, Quantification of the patterns of MEG activity for warm colors within one luminance level were more similar than patterns of MEG activity for cool colors within one luminance level. Shading shows the 95% C.I. Similarity was determined using the representational dissimilarity matricies at each time point relative to peak decoding accuracy (the RDM corresponding to the peak decoding time is shown in Figure 2e). At peak of the curve, p=0.014; total number of significant 5 ms bins: 33; longest span of significance: 19 bins (25 ms to 120 ms); time point of maximum, - 35 ms. **c**, Quantification of the observation that patterns of MEG activity for hues of a given luminance level were, on average, more similar than patterns of activity for hues across luminance levels. At peak of the curve, p=5×10^−4^; total number of significant 5 ms bins: 71; longest span of significance: 60 bins (−60 ms to 240 ms); time point of maximum, −120 ms. other conventions as for panel (b). **d**, Quantification of observation that patterns of MEG activity elicited by warm colors across luminance levels were more dissimilar than those patterns elicited by cool colors across luminance levels. At peak of the curve, p=0.002; total number of significant 5 ms bins: 5; longest span of significance: 4 bins (−10 ms to 10 ms); time point of maximum, 0 ms. other conventions as for panel (b).

Panels b-d quantify the temporal dynamics of the three main similarity relationships. The shaded gray area indicates the 95% C.I, computed by bootstrapping over subjects. First, representations of the two warm colors were more similar than representations of the two cool colors within a luminance level (e.g., among light dots, the pink and orange dots are closer together than the blue and green dots; **Figure 3b**). Second, representations for hues of a given luminance contrast were more similar than representations for colors across luminance levels (light and dark hues are linearly separable), and this pattern existed for at least 200 ms after peak onset decoding (**Figure 3c**), a longer duration than the pattern shown in Figure 3b. Finally, light and dark versions of each of the cool colors were closer together compared to light and dark versions of warm colors (e.g., the two pink dots were further apart than the two blue dots), but this pattern was only significant for a brief window of time near peak decoding (**Figure 3d**).

### Distinct representations of colors and color words

Do color names by themselves conjure mental representations analogous to those elicited by colors? This possibility is not inconceivable by Stroop tests that show judgements about the color of text are relatively slow when the color of the text is incongruent with the color name spelled in the text (e.g. people tend to be slower judging the color of red text spelling the word “blue” than judging the color of blue text spelling “blue”). Thus one interpretation of the results in Figure 2f is that the universal patterns in color naming—warm colors associated with “light” and cool colors with “dark”; warm colors named with more precision than cool colors—arise because participants unavoidably conjure color names when they view the color spirals, and the mental representations of color names drive the pattern of MEG results to colored spirals. To test the extent to which neural representations of color instantiated by language and perception are comparable, we randomly interleaved trials with the words “green” and “blue”, written in achromatic text, and evaluated whether classifiers trained on the pattern of MEG activity elicited by the words could decode color from MEG activity elicited by green and blue spirals. We chose “green” and “blue” words because the corresponding colored stimuli showed the highest concordance between stimuli of the same hue but different luminance, so we reasoned that the pattern of MEG activity elicited by these words would have the greatest likelihood of predicting patterns of activity elicited by colored spirals. Of course, the brain might first generate a perceptual representation, then a cognitive representation that maps word labels onto this perception. So one might expect a delay between when the pattern of activity elicited by the colored spirals matches the pattern of activity elicited by the color words, which requires that we test for cross-temporal generalization of the MEG patterns.

Unsurprisingly, term identity could be readily decoded from the pattern of MEG activity (onset of 65 ms [95% CI: 65, 70]; peak accuracy of 0.7991% [0.7387, 0.8627] obtained at 100 ms [90, 120]; **Figure 4a**). The early onset of decoding accuracy is consistent with the sensitivity V1 to the difference in appearance of the two words. But classifiers trained on the MEG data obtained using the words could not decode the color of the spirals from MEG activity elicited by the blue and green spirals, at any time delay between training and testing (**Figure 4b** left panel). Similarly, classifiers trained on the MEG data obtained using the colored spirals could not decode the term identity from MEG activity elicited by words, at any time delay between training and testing (**Figure 4b** right panel). These results suggest that the earliest neural representations of color instantiated by language and perception are distinct. The results also imply that the pattern of MEG activity in response to colored stimuli shown in Figures 2 and 3 are not confounded by language; thus the analyses provide the first direct measure of the geometry of the neural representation of color space.

**Figure 4.**
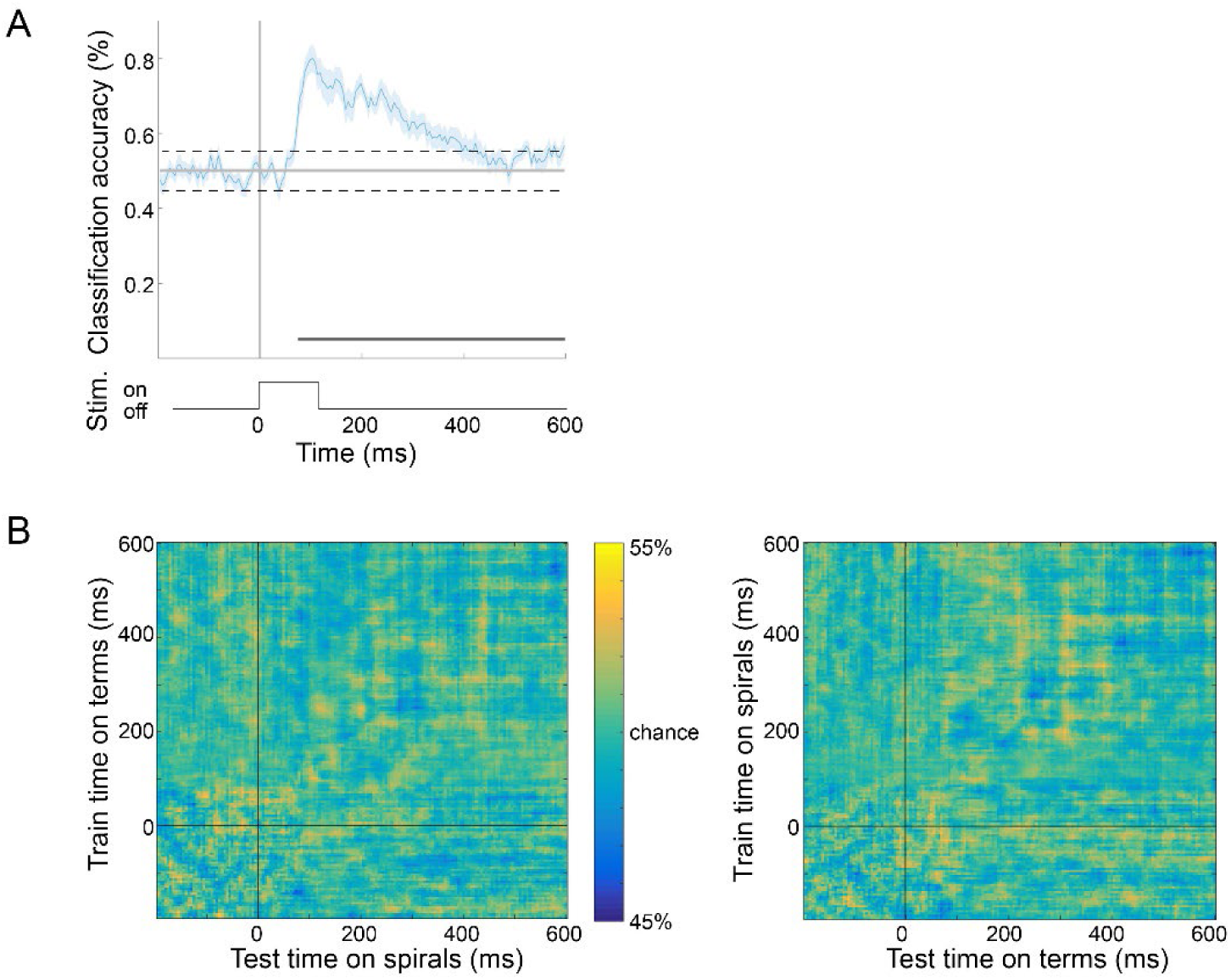
Spiral color cannot be decoded from MEG responses elicited by printed color names. **a**, Average accuracy of maxCorr classifiers (N=18 participants) decoding color term identity; the accuracy is likely attributed to slightly different retinotopic patterns of activation elicited by the two words. Plotting conventions as in Figure 2a. **b**, Cross-temporal decoding. Left: Average accuracy of maxCorr classifiers trained on color term data and tested on green and blue spiral data (accuracies averaged across both luminance levels of spirals). At every 5ms bin of the training data, a classifier was trained, and accuracy (color on the heatmap) was measured at every 5ms bin of test data. Right: maxCorr classifiers trained on green and blue spiral data and tested on color term data; same conventions as left.

Decoding was significant for a substantial amount of time, ranging from 55 to 590 ms (from Figure 2b). To what extent is the representation of color stable versus dynamic in time? The question can be addressed by examining the extent to which decoding patterns generalize across time [17, 29, 30]. Can classifiers trained at one point in time decode activity at another point in time? Several theoretical possibilities exist for the nature of the temporal representation. Two possibilities are the following. First, if the representation consists of a dynamic chain, with neural activity progressing through a series of unique states, classification performance will only be possible when test and train data are from the same time point. Second, if representations are relatively stable, it should be possible to classify activity obtained at one time point using training data obtained at a different time point. **Figure 5a** shows the cross-temporal generalization analysis for spiral identity, and **Figure 5b** indicates time points where decoding performance was significantly above chance. Decoding performance was strongest along the diagonal, consistent with the first possibility. But the plot also shows evidence of faint bands of significant decoding performance parallel to the diagonal, which is consistent with some version of the second possibility, specifically one in which a pattern of neural activity is reactivated at some later time point. To determine the time course of this reactivation, we reproduced the cross-temporal generalization analysis using 2 ms bins and performed a 2-D auto-correlation analysis on the temporal generalization plot. For the analysis, we restricted the train times to a window of 275-600ms. We shifted the temporal generalization plot horizotally along the test time axis and computed the 2-D Pearson’s correlation coefficient between the original plot and each of the shifted plots. We computed the second derivatives of the correlation coefficients and performed a Fourier analysis on the resulting time series. The frequency with the maximum amplitude was 20 Hz, which corresponds to a reactivation time of 50 ms (**Figure 5c)**.

**Figure 5.**
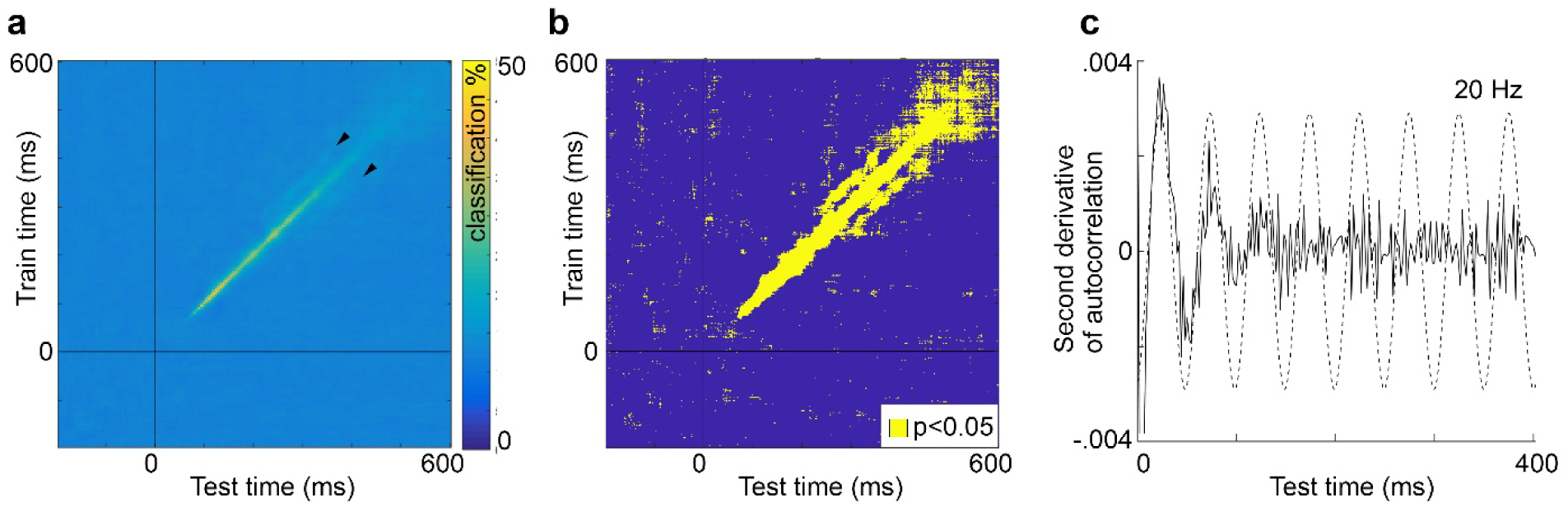
Cross-temporal generalization of color identity. **a**, Mean classification accuracy across participants (N=18) in classifying the eight color spirals. In addition to the strong classification accuracy along the diagonal, the plot shows faint bands of significant classification accuracy parallel to the diagonal (arrowheads). **b**, Plot showing time points of significant decoding, estimated by bootstrapping (p<0.05). **c**, Analysis of the spatial pattern in panel (a) to determine the temporal profile of the parallel bands. We performed a 2-D autocorrelation analysis on cross-temporal decoding accuracies by shifting the cross-temporal decoding map horizontally along the test-time axis and restricting the train-time axis (2 ms bins; 275 ms and 600 ms windows for the train-time and test-time axes respectively). We computed the 2-D Pearson’s correlation coefficient of the original map with each of the shifted maps and performed a Fourier analysis on the second derivatives of the 2-D correlation coefficients. The plot shows the second derivatives with a sine wave that has a frequency equal to the peak of the single-sided amplitude spectrum. The inset shows the single sided amplitude spectrum of the second derivative time series; shading shows the 99.95% CI of the scrambled second derivatives plot.

## Discussion

This paper presents an approach for investigating the similarity relationships among colors using multivariate analyses of MEG responses. The MEG data were obtained using colored stimuli defined by the cone-opponent cardinal mechanisms thought to be implemented by bipolar cells of the retina. MEG reflects predominantly cortical activity; thus any asymmetries in the MEG responses provide clues to the transformation by the cortex of the subcortical representation of color. Colors could not be decoded from patterns of MEG activity elicited in response to presentation of color words, which suggests that the decoding of color stimuli does not arise from implicit linguistic associations or implicit category effects of colors and words. Instead we interpret the results as evidence of the geometry of the neural representation of color, which, we suggest, is what accounts for perceptual similarity judgements that drive many universal patterns in color naming.

The results uncovered three asymmetries in the cortical representation of color related to color-naming patterns. One asymmetry reflects an interaction of luminance contrast and hue along the warm-cool axis of color space: warm colors were better decoded when they were relatively high luminance contrast, while cool colors were better decoded when they were relatively low luminance contrast. This asymmetry reflects a universal pattern in color naming in which “light” and “dark” are associated with “warm” and “cool” [23-25]. A second asymmetry shows that the neural representation elicited by stimuli of the same hue across luminance levels were more dissimilar for warm colors compared to cool colors. This pattern of results correlated with the extent to which participants used the same term for both light and dark versions of each hue. For example, the patters of MEG activity elicited in response to light and dark pink were more dissimilar than the patterns of MEG activity elicited in response to light and dark green; and participants are more likely to use different terms for the light and dark versions of pink than they are to use different terms for the light and dark versions of green. The MEG results show that patterns of activity elicited warm colors were more precisely encoded, and therefore more accurately decoded compared to cool colors, which may account for the greater naming efficiency of warm colors (Gibson et al, 2017). A third asymmetry reflects the preeminence of red among all colors. Regardless of luminance contrast, pink was decoded with the highest decoding accuracy and with the shortest onset latency. This result cannot be attributed to differences in contrast of the stimuli (the stimuli were all matched in cone contrast). The result echoes observations in monkeys showing that reddish hues produce very high gamma oscillations [31], and provides a neural correlate for the preeminence of red among all chromatic colors in color naming [23-25].

In addition to the three asymmetries described above that relate to universal patterns in color naming, the results uncover two other asymmetries. First, classification was slightly worse for colors corresponding to the daylight locus (orange-blue) compared to the anti-daylight locus (pink-green). This observation may have implications for understanding neural mechanisms of color constancy [32-36]. Second, the neural representations for colors at a given luminance level were more similar between the two warm colors than between the two cool colors. How does this surprising result relate to behavior? Color naming studies have shown that across luminance levels, warm colors are more readily distinguished compared to cool colors (see **Figure 1b**). But to our knowledge, color naming studies have not shown (or explicitly tested) whether colors at a given luminance level are less efficiently communicated for warm colors compared to cool colors. We wonder whether the result reflects the adaptation of the visual cortex to the statistics of the colors of objects, which are biased towards having warm colors [37]. Regardless, the set of results underscore the existence of a representation of hue that cannot be completely decoupled from a representation of luminance contrast. The existence of this neural representation may explain why establishing a uniform representation of color, in which hue and luminance contrast constitute independent dimensions, has been so difficult. Indeed, the present results suggest that the pursuit of a perceptually uniform color space may be ill founded, even if it has intuitive appeal.

The plots of cross-temporal generalization show a pattern of bands parallel to the identity diagonal which provides a rare empirical example (to our knowledge the first) of a theoretical possibility—the reactivation pattern, described by King and Dehaene [17]. This pattern suggests that color is encoded by a unique sequence of brain states that is reactivated at some later time point. The reactivation time course evident in this plot is 50 ms, implying that the brain state encoding color information depends on recurrent activation loops of 20 Hz. We speculate that these reactivation loops involve the hierarchical network of color-biased regions within the ventral visual pathway [38-40], that connect primary visual cortex with frontal cortex [41, 42].

In sum, despite what seems like perennial failure, the quest to identify a uniform representation of color persists. But the fact remains: the underlying principles that govern the geometric relationships among colors—the neural activity that clinical exams such as the Farnsworth hue-sorting test purportedly assess—and the connection between perceptual representations and color naming, remain unknown [16, 27, 43]. Indeed, the existence of so many color spaces exposes a deep mystery about how color is encoded by the brain. The present work establishes a proof of concept, using multivariate analysis of MEG activity to recover the similarity relationships among colors. A next step will be to measure responses to more colors, and to colors that vary in saturation, to establish more finely how color space is represented in the brain. In future work, we aim to develop of a complete color solid in which the spacing of the colors corresponds to equal distances in the dissimilarity of the neural responses.

## Methods

### Visual Stimuli

#### Visual Stimuli

Stimuli were eight square-wave spiral gratings on a neutral gray background (**Figure 1a**) [10, 44, 45] or achromatic words (white text on a gray background) of the words “green” and “blue”. The color words “green” and “blue” were chosen for the experiment because they were the terms used with the highest consensus in the color-naming experiment. Participants were asked to imagine the corresponding spiral color indicated by the term each time they saw the words. The 8 stimulus colors, four hues at two luminance-contrast levels, of matched cone contrast, were defined in DKL color space [19, 20] using implementations by Westland [46] and Brainard [47]: the axes of this color space are defined in terms of activation of the two cone-opponent post-receptoral chromatic mechanisms (**Figure 1a**). The z-axis is defined by luminance. The four hues were defined by the intermediate axes of DKL space: at 45° (pink), 135° (blue), 225° (green), and 315° (orange). Two spirals – one high luminance (20° elevation; “light”) and one low luminance (340° elevation; “dark”) -- were created at each hue. The neutral adapting background was 50 cd/m2. The luminance contrast of the stimuli was 26%. Modulation of the cone-opponent mechanisms, shown in **Figure 1a**, was computed relative to the adapting background gray, using the Stockman and Sharpe 2degree cone fundamentals, Judd corrected.

### Color-naming

Color names for the eight spirals used in the MEG experiments were obtained from fifty-two participants tested online, and the eighteen participants from the MEG experiment, who were tested in laboratory using the same display used for the MEG experiments.

The 52 online participants (29 female, age 19-63 years) were recruited and tested using Amazon Mechanical Turk (mTurk) and provided monetary compensation for their participation. They were shown each of the color stimuli (**Figure 1**) three times in random order and given the prompt “The color of the spiral is. (Use one word that you think other people would use to describe this spiral).” Only one response per spiral per participant was used in the analysis (selected at random from among the three responses per stimulus provided by each participant). The spiral was presented on a neutral gray background, as large as the screen would allow. Before beginning the experiment, participants were presented with a consent screen notifying them that participation was anonymous and that the task could be completed only once. In addition to color answers, some basic demographic information was collected (age, gender, language spoken, nationality, education level, vision, colorblindness, normal sleep/wake times, and handedness). Participants were excluded if English was not their first language or if they reported color blindness.

The 18 MEG participants completed the color-naming task during the first of two MEG sessions. Participants sat in the unlit MEG room before performing the main experiment and were presented with each stimulus three times in random order. Upon each presentation, participants were instructed to speak a one-word name for the color, which was relayed to the researcher sitting outside the room via 2-way intercom. Participants controlled the duration they saw each spiral via button press.

The results from the two sets of participants (online vs in-lab) were comparable and consistent with predictions based on the literature. For (MTurk|in-lab) participants, the same term was used at both luminance levels for hue 315° by (31%|61%); for hue 45° by (56%|56%); for hue 135° by (83%|94%); and for hue 225° by (85%|100%). The percentage of people who used the same term for the light and dark versions was the same for the MTurk and in-lab participants (Chi-square test of proportions, Bonferroni corrected for multiple comparisons alpha = 0.0125; for hue 45°, p =0.99; for hue 135°, p =0.22; for hue 225°, p =0.08; for hue 315°, p =0.02). These results showed that participants in both experiments tended to use the same label for the high and low luminance version of each hue more often when labeling cool colors (hues 135° and 225°) than warm colors (hues 45° and 315°). The most commonly used terms for hues 135° and 225° were the same for the MTurk participants and the in-lab participants (blue, green; consistent across luminance contrasts). The most commonly used terms for hue 45° were pink|pink (light|dark MTurk participants) and pink|purple (in-lab); for hue 315°, yellow|yellow (MTurk); orange|brown (in-lab). Because the results were comparable between the two sets of participants, we combined the data in the analyses that relate the MEG results to color naming.

### MEG Acquisition and Preprocessing

Participants were scanned in the Athinoula A. Martinos Imaging Center of the McGovern Institute for Brain Research at the Massachusetts Institute of Technology (MIT) over the course of 2 sessions, on an Elekta Triux system (306-channel probe unit consisting of 102 sensor triplets, with 204 planar gradiometer sensors, and 102 magnetometer sensors). Stimuli were back-projected onto a 44” screen using a SXGA+ 10000 Panasonic DLP Projector, Model No. PT-D10000U (50/60Hz, 120V). Data was recorded at a sampling rate of 1000Hz, filtered between 0.03-330Hz. Head location was recorded by means of 5 head position indicator (HPI) coils placed across the forehead and behind the ears. Before the MEG experiment began, 3 anatomical landmarks (bilateral preauricular points and the nasion) were registered with respect to the HPI coils, using a 3D digitizer (Fastrak, Polhemus, Colchester, Vermont, USA). During recording, pupil diameter and eye position data were collected simultaneously using an Eyelink 1000 Plus eye tracker (SR Research, Ontario, Canada) with fiber optic camera.

Once collected, raw data was preprocessed to offset head movements and reduce noise by means of spatiotemporal filters (Taulu et al, 2004; Taulu & Simola, 2006), with Maxfilter software (Elekta, Stockholm). Default parameters were used: harmonic expansion origin in head frame = [0 0 40] mm; expansion limit for internal multipole base = 8; expansion limit for external multipole base = 3; bad channels omitted from harmonic expansions = 7 s.d. above average; temporal correlation limit = 0.98; buffer length = 10 s). In this process, a spatial filter was applied to separate the signal data from noise sources occurring outside the helmet, then a temporal filter was applied to exclude any signal data highly correlated with noise data over time. Following this, Brainstorm software (Tadel et al., 2011) was used to extract the peri-stimulus MEG data for each trial (−200 to 600 ms around stimulus onset) and to remove the baseline mean.

### MEG Participants and Task

All participants (N=18, 11 female, age 19-37 years) had normal or corrected-to-normal vision, were right handed, and spoke English as a first language. One participant was an author and thus not naïve to the purpose of the study. During participants’ first session, they were screened for colorblindness using Ishihara plates; they also completed a version of a color-naming task as part of a separate study. After this task, participants completed a 100-trial practice session of the 1-back task that would be used in the MEG experimental sessions. Once this was complete, participants were asked if they had any questions about the task or the experiment; eye-tracking calibration was performed; and MEG data collection began.

During stimulus presentation, participants were instructed to fixate at the center of the screen. Spirals were presented subtending 10° of visual angle, for 116 ms, centered on the fixation point, which was a white circle that appeared during inter-trial intervals (ITIs, 1s). In addition to the spirals, the words “green” and “blue” were presented in white on the screen for the same duration, and probe trials to evaluate task performance were presented with a white “?”. Following each probe trial, which occurred every 3-5 stimulus trials (pseudorandomly interspersed, 24 per run), participants were instructed to report via button press if the two preceding spirals did or did not match in color (1-back hue task). Maximum response time was 1.8s, but the trials advanced as soon as participants answered.

Participants were encouraged to blink only during probe trials, as blinking generates large electrical artifacts picked up by the MEG. Each run comprised 100 stimulus presentations, and participants completed 25 runs per session over the course of approximately 1.5 hours. Between each run, participants were given a break to rest their eyes and speak with the researcher if necessary. Once 10s had elapsed, participants chose freely when to end their break by button-press. Over the course of both sessions, participants viewed each stimulus 500 times. Data from all participants was used (no data was excluded because of poor behavioral performance).

In addition to the 18 participants analyzed in the main thrust of this study, two pilot versions of the experiment were deployed. The first, more limited pilot experiment, was deployed with 2 participants (1 female, age 20-30 years) to determine the behavioral task and decoding parameters. The data from these participants was used to choose the parameters for the decoding analysis used in the rest of the study (see below). The second pilot experiment was more extensive, involving four participants, four colors, and 500 stimulus presentations per stimulus. This experiment was used to evaluate the power for color decoding; the results of the second pilot experiment were presented at the Society for Neurosciences Annual meeting in 2016 [12].

All experimental procedures involving participants tested in laboratory were approved by the Wellesley College Institutional Review Boards, the Massachusetts Institute of Technology Committee on the Use of Humans as Experimental Subjects, and the National Institutes of Health Intramural Institute Clinical Research Review Committee.

### MEG Processing and Decoding Analyses

Brainstorm software was used to process MEG data. Trials were discarded if they contained eyeblink artifacts, or contained out-of-range activity in any of the sensors (0.1-8000 fT). Three participants exhibited sensor activity consistently out of range, so this metric was not applied to their data as it was not a good marker of abnormal trials. After excluding bad trials, there were at least 375 good trials for every stimulus type for every participant. Data were subsampled as needed to ensure the same number of trials per condition were used in the analysis.

Decoding was performed using the Neural Decoding Toolbox (NDT) [48]. We used the maximum correlation coefficient classifier in the NDT to train classifiers to associate patterns of MEG activity across the sensors with the visual stimuli presented. This classifier computes the mean population vector for sets of trials belonging to each class in the training dataand calculates the Pearson’s correlation coefficient between those vectors and the test points. The class with the highest correlation is the classifier’s prediction. The main conclusions were replicated when using linear support vector machine classifiers. The classifiers were tested using held-out data— i.e. data that was not used in training. Data from both magnetometers and gradiometers were used in the analysis, and data for each sensor was averaged into 5-ms non-overlapping bins from 200 ms before stimulus onset to 600 ms after stimulus onset.

Custom MATLAB code was used to format MEG data preprocessed in Brainstorm for use in the NDT and to combine the two data-collection sessions for each participant. Decoding was performed independently for each participant, and at each time point. For each decoding problem, at each timepoint (a 5 ms time bin), the 375 trials for each stimulus condition were divided into 5 sets of 75 trials. Within each set, the 75 trials were averaged together. This process generated 5 cross-validation splits: the classifier was trained on four of these sets, and tested on one of them, and the procedure was repeated five times so that each set was the test set once. This entire procedure was repeated 50 times, and decoding accuracies reported are the average accuracies across these 50 decoding “runs”. This procedure ensured that the same data was never used for both training and testing, and it also ensured the same number of trials was used for every decoding problem. The details of the cross-validation procedure, such as the number of cross-validation splits, were determined during the pilot experiments to be those that yielded a high signal-to-noise ratio (SNR) and high decoding accuracy in both participants on the stimulus identity problem.

On each run, both the training and test data were z-scored using the mean and standard deviation over all time of the training data. Following others, we adopted a de-noising method that involved selecting for analysis data from the most informative sensors [49]; we chose the 25 sensors in the training data whose activity co-varied most significantly with the training labels. These sensors were identified as those with the lowest p-values from an F-test generated through an analysis of variance (ANOVA); the same sensors were then used for both training and testing. The sensor selection was specific for each participant. The sensors chosen tended to be at the back of the head. Analyses using all channels, rather than selecting only 25, yielded similar results.

Chance classification performance was 1/8 for all analyses except when comparing classification performance using data elicited by hues versus words, in which case classification was binary (chance equal to 50%). For each problem, a classifier was trained and tested in 5ms bins from time t=200ms before stimulus onset to t=600ms after stimulus onset. The classifiers’ performances were generated through a bootstrapping procedure. First, the problems were evaluated for each participant (resulting in 18 independent decoding time courses). Then, for each unique problem, we averaged the decoding time courses across participants. The gray shading in Figure 2a shows the standard error of the bootstrap mean. Significant decoding was defined as time bins in which decoding accuracy was above chance for five consecutive 5ms time bins. For the cross temporal generalization analysis, each classifier trained using data obtained at each time bin was tested using data obtained at every 5 ms time bin from −200 to 600 ms after stimulus onset creating a 2-dimensional matrix of decoding results.

Decoding analyses were also performed using eye tracking data collected during the MEG sessions. Two analyses were conducted: one using pupil diameter and one using eye position (**Figure S2)**. All parameters were identical to the MEG analysis except for the number of input features to the classifier. Rather than MEG sensors, the classifier used either the diameters of the two pupils (two features) or the xy coordinates of the positions of the two eyes (four features).

### Representational Similarity Analysis and Multidimensional Scaling Analysis

To examine the similarities between neural representations of colors, we obtained the results using ‘forced-error’ classifier tests. In this analysis, classifiers were trained and tested as in the other analyses, but the classifier was prevented from choosing the true stimulus. For each problem, the classifier returned the stimulus that elicited the most similar pattern of neural activity as elicited by the correct stimulus. This approach was designed not only to yield more data about dissimilarities between stimuli (which could be obtained just by analyzing the errors in the classifier), but also because it is directly analogous to the asymmetric matching task used in behavioral experiments aimed at recovering perceptual similarity among different colors. The dissimilarity matrix was then used in multidimensional scaling analysis, to uncover the geometry of the neural representation projected onto two dimensions. First, we constructed a confusion matrix of the errors the classifier made for each stimulus, for each individual participant (each column of which was normalized to sum to 1). Then, this matrix was averaged across the diagonal (for instance the instances when light pink was mistaken for light orange were averaged with the instances when light orange was mistaken for light pink) and normalized within each subject so that the largest percentage was equal to 1 and the smallest equal to 0. Finally, these values (measures of confusion) were subtracted from 1 to yield a representational dissimilarity matrix for each participant. These RDMs were averaged together and normalized 0-1.

## Notes

### Competing Interest Statement

The authors have declared no competing interest.

